# The *Ustilago hordei*-barley interaction is a versatile system to characterize fungal effectors

**DOI:** 10.1101/2020.12.10.419150

**Authors:** Bilal Ökmen, Daniela Schwammbach, Guus Bakkeren, Ulla Neumann, Gunther Doehlemann

## Abstract

Obligate biotrophic fungal pathogens, such as *Blumeria graminis* and *Puccinia graminis*, are amongst the most devastating plant pathogens, causing dramatic yield losses in many economically important crops worldwide. However, a lack of reliable tools for the efficient genetic transformation has hampered studies into the molecular basis of their virulence/pathogenicity. In this study, we present the *U. hordei*-barley pathosystem as a model to characterize effectors from different plant pathogenic fungi. We have generated *U. hordei* solopathogenic strains, which form infectious filaments without presence of compatible mating partner. Solopathogenic strains are suitable as heterologous expression system for fungal virulence factors. A highly efficient Crispr/Cas9 gene editing system is made available for *U. hordei*. In addition, *U. hordei* infection structures during barley colonization were analyzed by transmission electron microscopy, which shows that *U. hordei* forms intracellular infection structures sharing high similarity to haustoria formed by obligate rust and powdery mildew fungi. Thus, *U. hordei* has high potential as a fungal expression platform for functional studies of heterologous effector proteins in barley.

## Introduction

Plant pathogens have evolved different types of pathogenic life-styles with their host, ranging from obligate biotrophic, biotrophic, hemibiotrophic and necrotrophic. For successful colonization, each pathogen deploys a distinct set of effectors that target specific host molecules, pathways and structures. Recent genome and transcriptome analyses of a wide range of phytopathogens provide new insights into the effector inventories of pathogens that have different pathogenic life styles [1–5]. It has been found that compared to biotrophic pathogens, necrotrophs and hemibiotrophs have more plant cell wall degrading enzymes, secondary metabolites and toxins in order to kill their host cells during the infection and feed on nutrients released from dead host cells [6–8]. On the other hand, effector catalogues of biotrophs appear to be more specialized, reflecting their ability to efficiently suppress host defenses including regulated cell death, since their survival strictly depends on living host cells [7, 8]. While many effectors from different facultative biotrophs, hemibiotrophs and necrotrophs have been functionally characterized, there is still limited mechanistic insight into effectors of obligate biotrophic filamentous pathogens. A main reason for this gap is the absence of efficient genetic transformation and gene deletion techniques to perform reverse genetics in obligate biotrophs.

Currently, functional characterization of effectors of obligate biotrophic pathogens is performed via different strategies. Effectors from these pathogens can be heterologously expressed *in planta* and their positive contribution to virulence can be determined via subsequent inoculation of these plants with several pathogens [9, 10]. However, heterologous expression of some effectors *in planta* can result in strong pleiotropic defects that compromise symptom evaluations. In another strategy, the type III secretion system (T3SS) of *Pseudomonas syringae* (for Arabidopsis) and *Pseudomonas fluorescens* or *Pseudomonas atropurpurea* (for wheat and barley) is used for functional characterization of several intracellular effectors from obligate biotrophs like rusts and powdery mildews [11–15]. Any growth promotion observed for *Pseudomonas* sp. transformants, which deliver the desired effectors into the host cell during infection, is interpreted as a positive contribution to virulence [11–15]. However, the T3SS of *Pseudomonas* sp. also has some drawbacks. For example, fungal effectors that require post-translational modifications for their activity will not be correctly produced by the *Pseudomonas* sp. system, since prokaryotes lack the molecular machinery necessary for these modifications. In addition, the T3SS system delivers effectors into host plant cells, and hence fungal effectors that play roles in the apoplast or are required for haustorium formation and function during host colonization, might not be identified. Furthermore, the function of some effectors from biotrophic pathogens is to avoid or suppress PAMP-triggered immunity (PTI) to promote disease establishment. PAMPs from bacterial and fungal pathogens are different because of their phylogenetic distance, so unless the signaling pathways that lead to PTI are not completely conserved, PTI responses induced by *Pseudomonas* sp. may not be evaded or suppressed by fungal effectors. In another method, to validate virulence function of obligate biotrophs’ effector genes during host colonization, a host-induced gene silencing (HIGS) assay was developed [16–18]. However, requirements of stable transgenic host lines of HIGS constructs make this method very laborious.

The non-obligate biotrophic fungal pathogen *Ustilago hordei* is a causal agent of covered smut disease on barley and oat plants. *U*. *hordei* belongs to the group of Ustilaginales, members of which infect many economically important crops including maize, wheat, barley, oat and sugar cane. Similar to other smut fungi, pathogenic development of *U. hordei* is coupled to sexual development [19]. For successful infection, two haploid sporidia of opposite mating types fuse to form an infectious dikaryotic filament that subsequently differentiate to form an appressorium, a swollen hyphal cell that leads to direct penetration of host epidermal cells. During plant colonization, *U. hordei* proliferates both extra- and intracellularly and forms haustorium-like feeding structures in the host cells [20]. The *U. hordei* reaches and establishes itself in the host meristem and then grows with the plant until floral meristem develops spikelets, which likely gives a cue to the fungus to multiply and sporulate. Massive proliferation and sporulation of the fugus in the barley inflorescence is displayed by mass production of dark brown smut teliospores [21].

*Blumeria graminis* f. sp. *hordei* (*Bgh*) and *Puccinia graminis* f. sp. *tritici* (*Pgt*) are obligate biotrophic pathogens which are the causal agents of powdery mildew and stem rust on barley, respectively [22, 23]. Unlike *U. hordei*, which can be cultured *in vitro*, both *Bgh* and *Pgt* have obligate biotrophic lifestyles and cannot be cultured outside the host. Therefore, the generation of stable fungal transformants is the main bottleneck to study these pathosystems at the molecular level. Despite their phylogenetic distance, *Bgh* (Ascomycota), *Pgt* and *U. hordei* (Basidiomycota) share significant similarities: they are barley or wheat pathogens, establish strictly biotrophic interactions with their host in which they form specialized intracellular feeding-structures, the haustoria [20, 22, 23]. These similarities prompted us to establish cell biological, molecular and genetic methods to use the *U. hordei*-barley as a model system for functional characterization of effector candidates from different filamentous phytopathogens.

## Material and Methods

### Plant and fungal materials

To isolate total genomic DNA from axenic culture, *Ustilago hordei* (4857-4) strain were incubated in YEPS_light_ (0.4% yeast extract, 0.4% peptone, and 2% saccharose) liquid medium at 22°C with 200 rpm shaking till OD:1.0. To isolate total gDNA from *U. hordei*-infected barley plants (at 6 dpi), the third leaves of the *U. hordei* infected barley plants were collected by cutting 1 cm below the injection needle sites. Leaf samples were then frozen in liquid nitrogen and ground using a mortar and pestle under constant liquid nitrogen. The gDNA was isolated by using a MasterPure™ Complete DNA&RNA Purification Kit (Epicentre^®^, Illumina^®^, Madison, Wisconsin, USA) according to the manufacturer’s instructions.

Susceptible Golden Promise barley cultivar was grown in a greenhouse at 70% relative humidity, at 22°C during the day and the night; with a light/dark regime of 15/9 hrs and 100 Watt m^−2^ supplemental light when the sunlight influx intensity was less than 150 Watt m^−2^.

### Nucleic acids methods

Fungal biomass quantification was performed by using quantitative PCR (qPCR) analysis as in Ökmen *et al*., 2018 [20]. Genomic DNA from infected barley leaves at 6 days post-inoculation (dpi) was isolated by using the MasterPure™ Complete DNA&RNA purification Kit (Epicentre®, Illumina®) according to manufacturer’s instructions. The *U. hordei UhPpi* gene (*UHOR_05685*) was used as a reference gene. A standard curve was constructed by using serial dilutions of *U. hordei* genomic DNA (100, 10, 1, 0.1, 0.01, 0.001 ng μl^−1^) using *UhPpi* as a reference gene. Base 10 logarithms of DNA concentrations were plotted against the crossing point of Ct values. The qPCR reaction was performed in a Bio-Rad iCycler system by using the following program: 2 min at 95°C followed by 45 cycles of 30 s at 95°C, 30 s at 61°C and 30 s 72°C. The primers that have been used for qPCR are listed in **Table S1.** All PCRs reactions were performed by using Phusion© DNA polymerase (Thermo Scientific; Bonn, Germany) following the manufacturer’s instructions with 100 ng genomic DNA or cDNA as template. All primers that are used in PCR reaction for cloning of different genes are listed in **Table S1**. The amplified DNA fragments were then used for cloning processes. All PCR reaction took place in a PTC-200 (Peletier Thermal Cycler, MJ Research) PCR machine. Nucleic acids deriving from PCRs or restriction digests reactions were purified from 1% TAE agarose gels with the “Wizard SV Gel and Purification System Kit” (Promega) according to the manufacturer’s instructions. Plasmid isolation from bacterial cells was performed by using the QIAprep Mini Plasmid Prep Kit according to the manufacturer’s information.

### Construction of expression vectors

For heterologous gene expression constructs (*p123-pUHOR02700::SP-Gus-mCherry*, *p123-pUHOR02700::Gus-mCherry*, *p123-pUHOR02700::SP-FvRibo1* and *p123-pActin::SP-CfAvr4*), standard molecular biology methods were used according to molecular cloning laboratory manual of Sambrook *et al*. (1989) [24]. Amplified PCR fragments for each gene (*Gus-mCherry*, *FvRibo1* and *CfAvr4*) were cut with appropriate restriction enzymes, subsequently they were ligated into a vector that was digested with the same restriction enzymes by using T4-DNA ligase (New England Biolabs; Frankfurt a.M., Germany) according to manufacturer’s instructions. The sequences confirmation of each construct was performed via sequencing at the Eurofins Genomics (Cologne, Germany). All vector constructs, primer pairs and restriction sites are indicated in **Table S1.** *Escherichia coli* transformation was performed via heat shock assay according to standard molecular biology methods [24].

### CRISPR/Cas9 gene editing system

To establish the CRISPR/Cas9-HF (high fidelity) gene editing system in the *U. hordei*, a plasmid containing codon optimized *Cas9*-*HF* gene under the control of *Hsp70* promoter and carboxin resistance was used from Zuo et al. (2020) [25]. To express sgRNA for targeted gene, *Ustilago maydis pU6* promotor was replaced with the *U. hordei pU6* promotor. The sgRNAs for the knockout of the *U. hordei* gene was designed by E-CRISPR (http://www.e-crisp.org/ECRISP/aboutpage.html) **(Table S1)** [26]. Plasmid construction for CRISPR/Cas9 was performed as described by Zou *et al*. (2020). The CRISPR/Cas9-HF vector was linearized with restriction enzyme Acc65I, and subsequently assembled with spacer oligo and scaffold RNA fragment with 3’ downstream 20 bp overlap to the plasmid by using Gibson Assembly [27].

### Fungal transformation and virulence assays

The *U. hordei* transformation assay was conducted by using protoplasts according to Kämper, (2004) [28]. Virulence assays for DS200-FvRibo1 and DS200 *U. hordei* strains were performed according to Ökmen *et al*., 2018. Briefly, all *U. hordei* strains were grown in YEPS_light_ liquid medium at 22°C and 200 rpm shaking until getting an OD_600_ of 0.6-0.8. Subsequently, *U. hordei* cells were centrifuged at 3500 rpm for 10 min at RT and resuspended in sterile distilled water supplemented with 0.1% Tween-20 to an OD_600_ of 3.0. Then each *U. hordei* cell suspension was injected into stems of 12-day-old barley seedlings (Golden Promise) with a syringe with needle. All infection assays were performed in three biological replicates with at least 15 plants. Fungal biomass quantification of *U. hordei* was performed at 6 dpi by using genomic DNA (200 ng μl^−1^) as the template with qPCR. To confirm secretion of GUS-mCherry protein in apoplast of barley leaf, apoplastic fluid (AF) from *U. hordei* DS200 ±GUS-mCherry strains infected barley leaves were isolated according to van der Linde *et al*. (2012) [29]. After AF isolation, western blot analysis was performed to detect GUS-mCherry signal in isolated AFs. Western blot was performed as described in Mueller *et al.* (2013) [30]. To confirm that *U. hordei* can express and secrete functional effector proteins from different fungi, *Avr4* of *Cladosporium fulvum* and *Ribo1* (encoding a secreted ribotoxin) of *Fusarium verticillioides* were heterologously expressed in the *U. hordei* strain DS200 with the *UHOR_02700* signal peptide (SP) and under control of *pActin* (for *in vitro* expression) and *pUHOR_02700* (for *in planta* expression) promoters, respectively. To confirm expression and secretion of CfAvr4 effector protein in *U. hordei in vitro*, culture filtrates isolated from DS200-CfAvr4, DS200-FvRibo1 and DS200 strains (OD:1.0) were collected with centrifugation (13000 rpm for 5 min). After filter sterilization, each culture filtrate was infiltrated in tobacco leaves expressing *Cf4* resistance gene (a gene encoding tomato Cf4 receptor protein that can recognize CfAvr4 protein and induce hypersensitive response) to induce hypersensitive response.

### Light Microscopy

The wheat germ agglutinin (WGA)-AF488 (Molecular Probes, Karlsruhe, Germany) and propidium iodide (PI) (Sigma-Aldrich) staining was performed according to Ökmen *et al*., 2018 [20]; WGA-AF488 stains fungal cell walls and the propidium iodide stains plant cell walls. *U. hordei* infected barley leaves were first bleached in pure ethanol and then they were boiled for 1-2 hours in 10% KOH at 85°C. Subsequently, the pH of boiled leaf samples was neutralized by using 1xPBS buffer (pH: 7.4) with several washing steps. Then, the WGA-AF488/PI staining solution (1 μg ml^−1^ propidium iodide, 10 μg ml^−1^ WGA-AF488; 0.02% Tween 20 in PBS pH 7.4) was vacuum infiltrated in samples for 5 min at 250 mbar using a desiccator (vacuum infiltration step was performed three times). The WGA-AF488/PI stained leaf samples were stored in 1xPBS buffer (pH: 7.4) at 4°C in the dark until microscopy. WGA-AF488: excitation at 488 nm and detection at 500-540 nm. PI: excitation at 561 nm and detection at 580–630 nm.

To visualize secretion of GUS-mCherry protein in *U. hordei* during barley colonization, *U. hordei* DS200 ±GUS-mCherry strains were inoculated on barley plants. Subsequently, infected barley leaves were checked for localization of GUS-mCherry at 4 dpi by using a Leica SP8 confocal microscopy. For mCherry fluorescence of hyphae in barley tissue, an excitation at 561 nm and detection at 580–630 nm was used.

### Transmission electron microscopy

Chemically fixed samples were prepared according to Wawra *et al*., (2019) with minor changes [31]. For TEM observation, 2 mm leaf discs from infected and non-infected *Hordeum vulgare* leaves were excised from 1 cm below infection sites by using a biopsy punch and chemically fixed in 2.5% glutaraldehyde and 2% paraformaldehyde in 0.05 M sodium cacodylate buffer, pH 6.9, supplemented with 0.025% CaCl2 (w/v) for 2 h at room temperature. Subsequently, samples were rinsed six times for 10 minutes in 0.05 M sodium cacodylate buffer (pH 6.9, rinse 3 and 4 supplemented with 0.05 M glycine) and post-fixed for 1 h at room temperature with 0.5% OsO4 in 0.05 M sodium cacodylate buffer, pH 6.9, supplemented with 0.15% potassium ferricyanide. After thorough rinsing in 0.05 M sodium cacodylate buffer (pH 6.9) and water, samples were dehydrated in an ethanol series from 10% to 100%, gradually transferred to acetone and embedded into Araldite 502/Embed 812 resin (EMS, catalog number 13940) using the ultra-rapid infiltration by centrifugation method revisited by McDonald (2014) [32].

For TEM observation, leaf samples were also processed by means of high pressure freezing and freeze substitution as an alternative to conventional chemical fixation following the procedure described in Micali *et al*. (2011) for ultrastructural observations [33]. Once the samples reached room temperature, they were rinsed in acetone, carefully removed from the aluminium specimen carriers and gradually infiltrated in LR White resin (Plano GmbH) for 6 days. Resin polymerization was done in flat embedding molds at 100°C for 24 h.

Ultrathin (70-90 nm) sections were collected on nickel slot grids as described by Moran and Rowley (1987) [34], stained with 0.1% potassium permanganate in 0.1 N H2SO4 [35], followed by 2% (w/v) aqueous uranyl acetate and lead citrate for 15 min [36] and examined with an Hitachi H-7650 TEM (Hitachi High-Technologies Europe GmbH, Krefeld, Germany) operating at 100 kV fitted with an AMT XR41-M digital camera (Advanced Microscopy Techniques, Danvers, USA). Immunogold labelling of ß-1,3-glucan was done according to the procedures described previously [33].

## Results

### Construction of solopathogenic strain of *Ustilago hordei*

The requirement of mating for the induction of pathogenic development implies that genetic mutations always need to be made in two compatible *U. hordei* strains, which presents an obvious drawback of the system, particularly for larger scale analyses. To optimize the work flow, we have generated a solopathogenic *U. hordei* strain, which does not require a mating partner to form an infectious filament **(Figure 1A-D).** For pathogenic development, *U. hordei* requires both a compatible pheromone (Mfa)/pheromone receptor (Pra) pair and an active heterodimer made from *bE* and *bW* gene products [37, 38]. To construct solopathogenic *U. hordei* strains, the *bE1* allele from mating type locus 1 (*MAT-1*) were replaced with the *bE2* allele from mating type locus 2 (*MAT-2*), or both *Mfa1* and *bE1* alleles from mating type locus 1 (*MAT-1*) were replaced with *Mfa2* and *bE2* alleles from mating type locus 2 (*MAT-2*). While the constructed DS199 solopathogenic strain had a compatible *b*-locus with *bE2* and *bW1* genes from different mating types, the DS200 solopathogenic strain contained both compatible MFA2/PRA1 and bE2/bW1 pairs to facilitate the formation of infectious filaments in the absence of a mating partner. For generation of DS199 and DS200 strains, homologous recombination constructs with an FRT-flanked hygromycin resistance cassette (for *Mfa2* construct) or phleomycin resistance cassette (for *bE2* construct), were used, which were removed from the genome after induction of the FRT recombinase [39].

**Figure 1.**
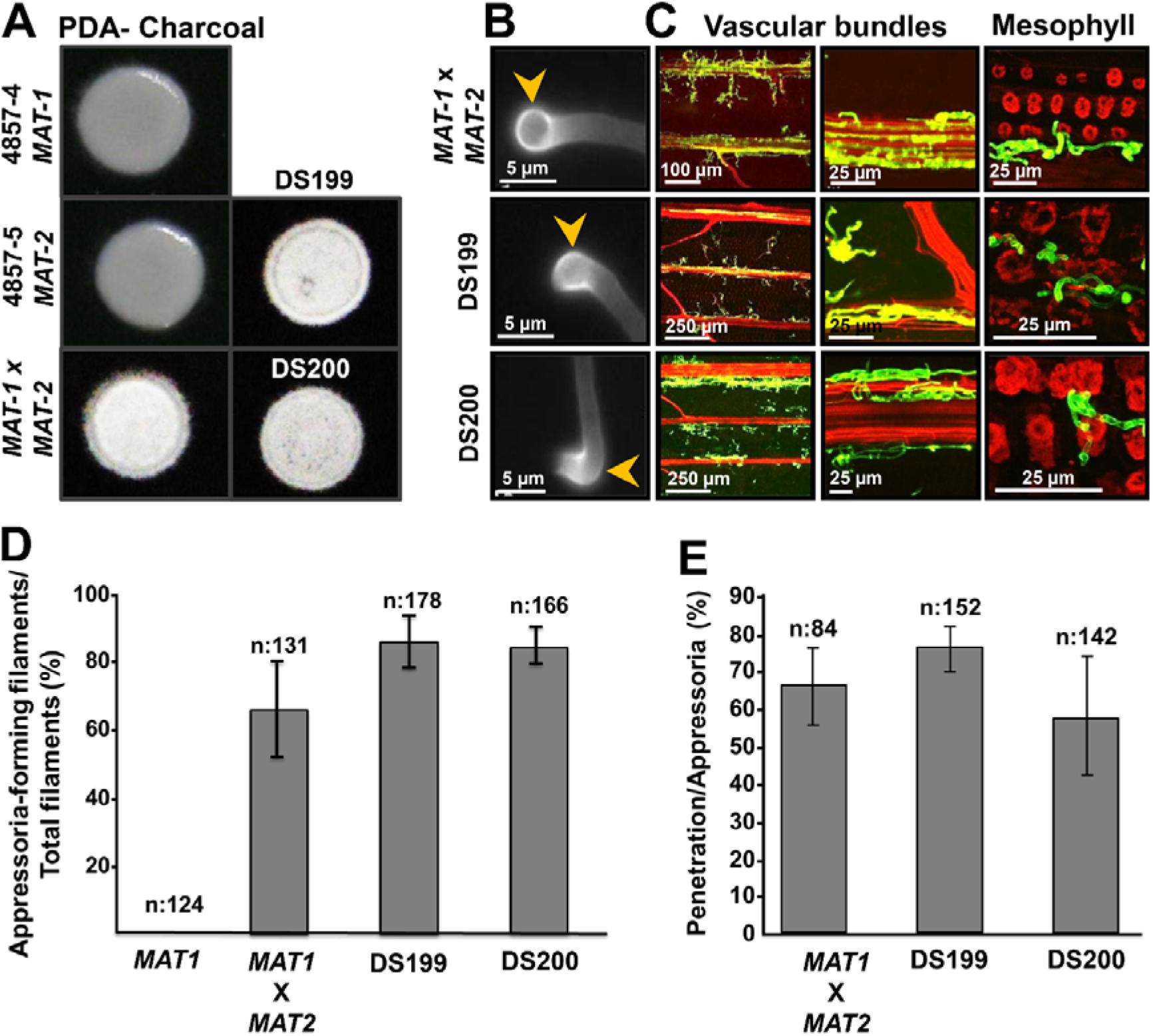
Generation of a solopathogenic *Ustilago hordei* strain. **(A)** Filamentation test on charcoal plate. *U. hordei* wild-type strains 4857-4 *MAT-1*, 4857-5 *MAT-2*, mating of 4857-4 *MAT-1*X4857-5 *MAT-2*, solopathogenic DS199 and DS200 strains. Pictures were taken after 3 days incubation at RT. **(B) Appressoria formation ability of** *Ustilago hordei* **strains on parafilm**. Mating of *U. hordei* wild-type 4857-4 *MAT-1* and 4857-5 *MAT-2*, solopathogenic DS199, solopathogenic DS200. Yellow arrowheads indicate appressoria. Pictures were taken after 24 hours incubation **(C) Disease development of different** *Ustilago hordei* **strains** *on* **barley.** Mating of *U. hordei* wild-type 4857-4 *MAT-1* and 4857-5 *MAT-2* at 3 dpi, solopathogenic DS199 at 3 dpi, solopathogenic DS200 at 3 days post inoculation (dpi). Following WGA-AF488/Propidium iodide staining, fungal cell walls are shown in green and plant cell walls in red. **(D) Quantification of appressoria formation for *U. hordei*** wild-type 4857-4 *MAT1*, solopathogenic DS199 and DS200 on plant. **(E) Quantification of penetration efficiency for *U. hordei*** wild-type 4857-4 *MAT1*, solopathogenic DS199 and DS200 on plant.

Both the DS199 and DS200 solopathogenic strains were then used for further analysis. To show filamentation ability of solopathogenic strains, single *U. hordei* 4857-4 *MAT-1* and 4857-5 *MAT-2* strains, mixed 4857-4 *MAT-1*X4857-5 *MAT-2*, and solopathogenic strains were grown on PD agar plates containing charcoal. While single 4857-4 *MAT-1* and 4857-5 *MAT-2*A strains were not filamentous, mated 4857-4 *MAT-1*X4857-5 *MAT-2* and solopathogenic strains were fully filamentous on PDA charcoal plates **(Figure 1A)**. To test whether the solopathogenic strains form infection structures (hyphal tip swellings; appressoria) that are required for host penetration, mated wild-type *U. hordei* 4857-4 *MAT-1* and 4857-5 *MAT-2*, and solopathogenic strains were sprayed on parafilm, which previously has been shown to artificially induce appressorium formation in *Ustilago maydis* [40]. This showed that both solopathogenic strains form appressoria that are comparable to wild-type strain **(Figure 1B)**. Quantification of appressoria formation on barley leaves revealed that there is no significant difference in appressoria formation **(Figure 1D)** and penetration efficiency **(Figure 1E)** of wild-type and solopathogenic strains. Wheat germ agglutinin-AF488/propidium iodide (WGA-AF488/PI) staining of wild-type and solopathogenic strains infecting barley leaves revealed that there is no visible difference in colonization at 3 days post-inoculation (dpi), i.e. all three strains were found to colonize the host mesophyll tissue **(Figure 1C)**. While the leaf colonization assay did not show obvious differences between solopathogenic and wild-type strains, in barley seed infection assays, the solopathogenic strain was rarely found to have colonized the barley inflorescence and produce teliospores (after 3-4 months post inoculation) **(Figure S1)**.

### Ultrastructure of the *Ustilago hordei-*barley interphase during biotrophic interaction

Similar to powdery mildews and rusts, smut fungi including *U. hordei* are considered to be intracellular pathogens. However, in all these interactions, the host plasma membrane is not breached and thus the host-pathogen interaction takes place through the biotrophic interphase, which consists of fungal cell wall (FCW), extracellular matrix (ECM) and plant cell wall (in some regions) **(Figure 2A-B)**. Both chemically fixed as well as high pressure-frozen samples were used to perform transmission electron microscopy (TEM) to display ultrastructural features of the *U. hordei-*barley biotrophic interphase during infection. Samples were taken at 8 dpi, when *U. hordei* growth is primarily intracellular and hyphae can be found in epidermal cells, mesophyll cells and in vascular bundles. Transmission electron micrographs show fungal hyphae growing inside plant cells and frequently branching **(Figure S2A-D)**. The fungal hyphae contain free ribosomes, strands of endoplasmic reticulum, mitochondria, nuclei, as well as lipid bodies, vesicles and vacuoles **(Figure S3A-F)**. Transmission electron micrographs also show the presence of closely paired nuclei in fugal hyphae, which are intimately associated with mitochondria **(Figure S3E)**. Vesicles and multivesicular bodies were also frequently observed at hyphal tips and in the plant cytoplasm adjacent to fungal penetration sites, respectively **(Figure 2C-D and Figure S3B, C, F)**.

**Figure 2.**
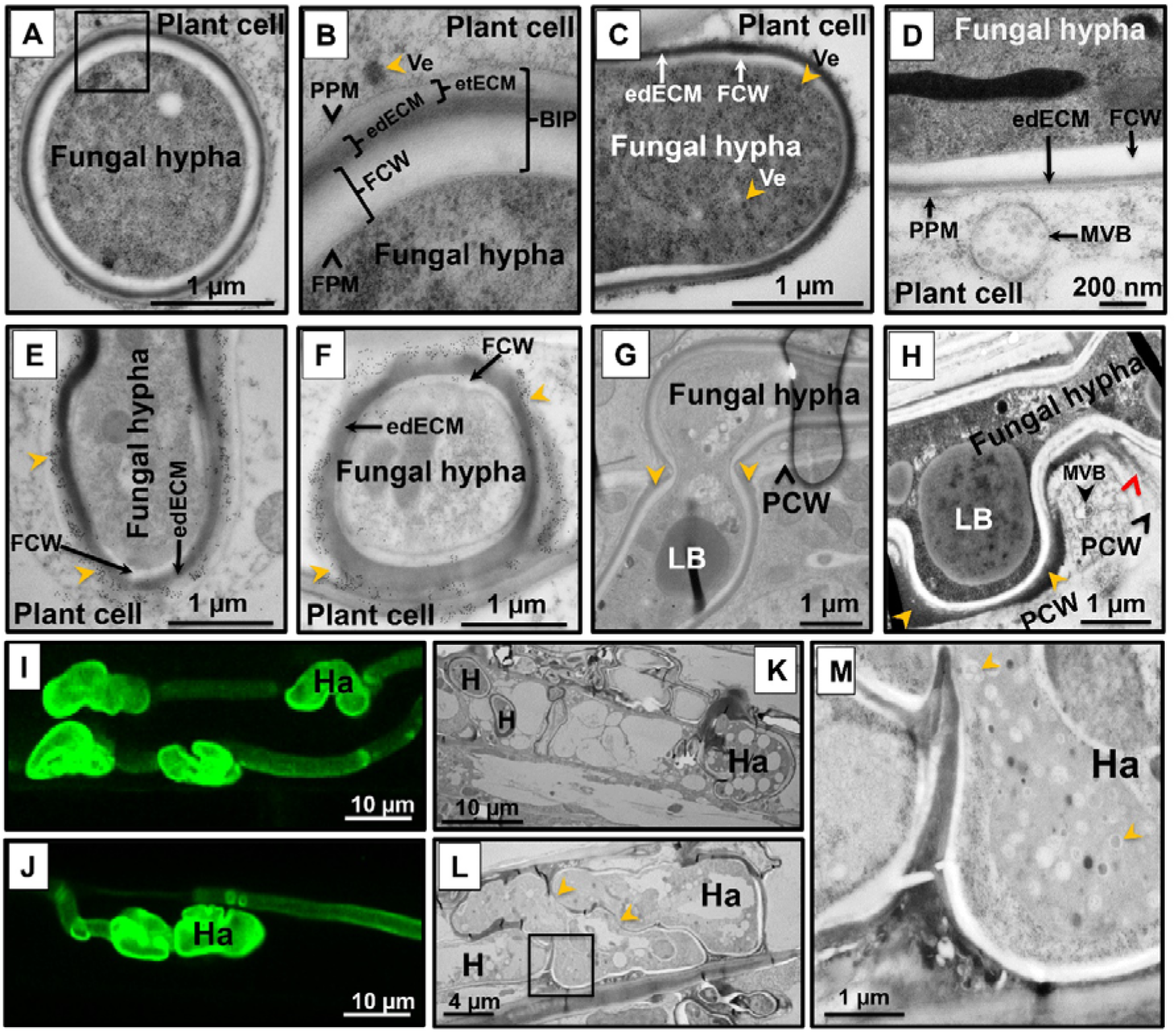
(A-H) Transmission electron microscopy micrographs of wild-type *Ustilago hordei* infected barley leaves. **(A, B) Biotrophic interphase in the** *U. hordei***-barley interaction.** During host colonization, *U. hordei* invaginates the host cell membrane without breaching it. The host-pathogen interaction mainly takes place within this biotrophic interphase (BIP), which consists of fungal cell wall (FCW), electron-dense extracellular matrix (edECM) and electron-translucent extracellular matrix (etECM). **(C, D) Formation of vesicles at the hyphal tip of** *U. hordei*. Fungal vesicles (Ve) with cores of different electron density and plant multivesicular bodies (MVB) were detected at hyphal tips and in the plant cytoplasm close to fungal penetration sites, respectively. **(E, F) Immunogold labeling of callose with a monoclonal antibody recognizing (1-3)-**β**-glucan epitopes.** Callose accumulation was detected at the electron-translucent ECM (IdECM) site (**yellow arrowheads)**. **(G, H) Cell-to-cell penetration of** *U. hordei*. *U. hordei* primarily grows intracellularly at 8 dpi in barley leaves. When the fungal hyphae penetrate a new plant cell, the hypha gets thickened at the site of cell-to-cell passage, resembling appressorial structures **(G)**. The edECM gets thicker where the hypha was in contact with the plant cell wall (yellow arrowheads) **(H)** and electron-dense material can also diffuse into adjacent part of the plant cell wall (red arrowhead) **(H)**. **(I-M) Haustoria formation during host colonization.** *U. hordei* grows intracellularly and forms haustorial structures in barley cells. **(I, J) WGA-AF488/Propidium iodide staining** was performed to visualize *U. hordei* at 8 dpi under confocal/ fluorescent microscopy. **(K-M) Transmission electron micrographs showing different planes of section through haustoria.** Haustorial structures were distinguished from hyphae by their bigger size and inter-connected lobular shapes. Yellow arrowheads **(L)** point out the connections between haustorial lobes. *U. hordei* haustoria possess large vacuoles with a granular lumen containing vesicles of different size **(M, yellow arrowheads; magnification of inset in L)**. BIP: Biotrophic Interphase; FCW: Fungal Cell Wall; FPM: Fungal Plasma Membrane; H: Hypha; Ha: Haustorium; edECM: electron-dense extracellular matrix; etECM: electron-translucent extracellular matrix; LB: Lipid bodies; MVB: Multi Vesicular Body; PCW: Plant Cell Wall; PPM: Plant Plasma Membrane; Ve: Vesicles.

It caught our attention that the *U. hordei* hyphal cell wall appears to be surrounded by a two layered extracellular matrix of different electron density: an inner electron-dense (edECM) and an outer electron-translucent layer (etECM), both of unknown composition **(Figure 2A-B, Figure S3A-C)**. Immunogold gold labeling with a monoclonal antibody specific for (1-3)-β-glucans showed that callose is present in the etECM **(Figure 2E-F)**. The edECM was particularly prominent (between 50-500 nm thickness) when the hyphae were in contact with the plant cell wall **(Figure 2H, yellow arrowheads, Figure S2A, D).** In some areas, the middle lamella of the plant cell wall close to the interface was more electron-dense than in areas where it was not in contact with the fungal hypha **(Figure 2H, red arrowhead)**. Another interesting observation of our TEM analysis was that at the site of cell-to-cell penetration, the *U. hordei* hypha is swollen, which resembles appressorial structures **(Figure 2G and Figure S2C)**. During barley colonization, *U. hordei* also forms structures similar to the haustoria known for obligate biotrophs, where they are described to function as feeding structures **(Figure 2I-M)**. Haustorial structures of *U. hordei* were distinguished from the normal hyphae by their bigger size and inter-connected lobular shapes **(Figure 2I-L)**. High magnification transmission electron micrographs of *U. hordei* haustoria showed that these structures possess large vacuoles with a granular lumen containing vesicles of different size **(Figure 2M)**.

### Heterologous gene expression in *Ustilago hordei*

To use the *U. hordei*-barley pathosystem for functional characterization of secreted virulence factors, heterologous gene expression was established in this smut fungus. As a proof of concept, *mCherry* fused to the *Escherichia coli GusA* gene under the control of the *U. hordei UHOR_02700* promoter (highly induced upon barley penetration) was heterologously expressed in the *ip* (*cbx*) locus of the solopathogenic DS200 strain, either with (+) or without (−) signal peptide (from *UHOR_02700*) for extracellular secretion **(Figure 3A)**. Confocal microscopy imaging was performed with DS200 strains expressing ±SP-GusA-mCherry on barley leaves at 3 dpi to monitor expression and localization of recombinant proteins. While SP-GusA-mCherry was localized around the hyphal tip region (showing secretion from the biotrophic hypha), -sp-GusA-mCherry was localized inside the fungal cytoplasm **(Figure 3A)**. Furthermore, western blot analysis was performed with apoplastic fluid isolated from barley leaves infected with ±SP-GusA-mCherry DS200 strains to confirm secretion of the recombinant proteins. Western blot results also showed that while the secreted full-length SP-GusA-mCherry (~100 kDa) and cleaved free mCherry (~27 kDa) were detected in isolated apoplastic fluid, the cytoplasmic-sp-GusA-mCherry was not detectable in isolated apoplastic fluid **(Figure 3B)**.

**Fig 3.**
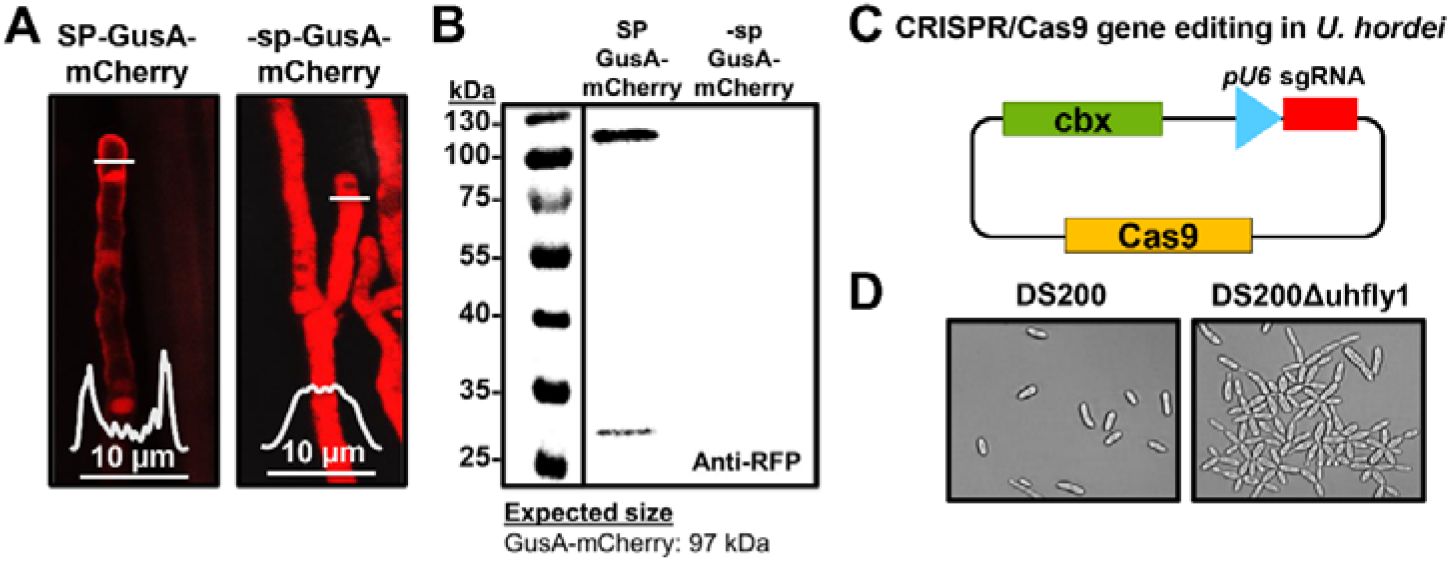
(A-B) Heterologous expression of *GusA-mCherry* in *Ustilago hordei*. **(A)** GusA-mCherry was heterologously expressed in solopathogenic strain DS200 under control of the *UHOR_02700* promotor with or without signal peptide for extracellular secretion. ±SP-GusA-mCherry DS200 strains were inoculated on barley seedlings and at 4 dpi confocal microscopy was performed to monitor expression and localization of recombinant proteins. While +SP-GusA-mCherry is secreted around the tip of the invasive hyphae, −sp-GusA-mCherry localizes in the fungal cytoplasm. The white graphs indicate the mCherry signal intensity along the diameter of the hyphae (illustrated by white lines in the image). **(B)** Western blot analysis was performed with apoplastic fluid isolated from barley leaves infected with ±SP-GusA-mCherry DS200 strains. While a band corresponding to secreted +SP-GusA-mCherry (at ~100 kDa) and free mCherry (at 27 kDa) in isolated apoplastic fluid was detected, no band corresponding to cytoplasmic −sp-GusA-mCherry could be detected. Anti-RFP antibody was used for western blot. **(C-D) Establishment of CRISPR/Cas9 gene editing system for *Ustilago hordei***. **(C)** Codon optimized *Cas9* was cloned into *p123* plasmid under the control of *Hsp70* promoter. The *U. hordei pU6* promotor was used to express sgRNA for the targeted gene. Carboxin resistance was used as selection marker. **(D)** *U. hordei Fly1* gene, a fungalysin metalloprotease involved in fungal cell separation, was edited via CRISPR/Cas9 system for knock-out. While DS200 sporidia showed normal growth, DS200Δuhfly1 cells were impaired in cell separation.

### CRISPR/Cas9 gene editing in *Ustilago hordei*

To establish the CRISPR/Cas9-HF (high fidelity) gene editing system in *U. hordei*, a codon optimized *Cas9*-*HF* gene under the control of *Hsp70* promoter was expressed in the solopathogenic strain DS200. The *U. hordei pU6* promotor was used to express sgRNA of a targeted gene **(Figure 3C)**. As a proof of concept, the *U. hordei Fly1* gene, a fungalysin metalloprotease involved in fungal cell separation in *U. maydis* [41], was edited via CRISPR/Cas9 system to result in a truncated protein (after aa 25, a stop codon was introduced). After sequence confirmation of the mutants, microscopic observations showed that while the DS200 strain formed normal yeast cells, DS200Δfly1 strains were impaired in cell separation in liquid medium indicating functional conservation of Fly1 in both *U. hordei* and *U. maydis* **(Figure 3D)**. 83.3 (±8.3)% selected independent transformed colonies showed an impaired cell separation phenotype, indicating high efficiency of this method in *U. hordei*.

### Activity of heterologous virulence factors expressed in *U. hordei*

To show that *U. hordei* can express and secrete functional proteins from different plant pathogenic fungi, *Avr4* from *Cladosporium fulvum* (a chitin-binding Avr protein that can be recognized by tomato resistance protein Cf4) and *Ribo1* (encoding a secreted ribotoxin) from *Fusarium verticillioides*, were heterologously expressed in the DS200 strain. For secretion from *U. hordei* hyphae, the open reading frames were fused with the sequence encoding *UHOR_02700* signal peptide (SP). For constitutive expression, heterologous genes were expressed under control of the *pActin* promoter, and for specific transcriptional induction during plant colonization, the promoter *pUHOR_02700* (highly expressed *in planta U. hordei* effector gene) was used [20]. To confirm *in vitro* expression and secretion of CfAvr4 effector protein in *U. hordei*, culture filtrates isolated from DS200-CfAvr4, DS200-FvRibo1 and DS200 strains were collected and infiltrated in *Nicotiana benthamiana* leaves expressing the *Cf4* resistance gene, a gene encoding the tomato Cf4 receptor protein which recognizes the CfAvr4 protein and induce hypersensitive response [42]. While neither DS200 nor DS200-FvRibo1 (expressed only *in planta*) culture filtrates did induce any hypersensitive response-mediated cell death in Cf4 expressing tobacco leaves, the culture filtrate of DS200-CfAvr4 induced hypersensitive response-mediated cell death in the presence Cf4 **(Figure 4A)**. Since Avr4-triggered HR cannot be observed in barley, we deployed FvRibo1 to test secretion of a functional heterologous virulence factor *in planta*. Heterologous expression of plant cytotoxic FvRibo1 protein in the DS200 strain was expected to negatively affect the growth of *U. hordei* on barley leaves. Macroscopic observations of infected barley leaves at 6 dpi showed that infection by DS200 strain causes spreading chlorosis along the leaf veins, reflecting spread of fungal proliferation **(Fig 4C)**. In contrast, DS200-FvRibo1 infected barley leaves displayed accumulated focal necrotic spots, reflecting restriction of fungal proliferation **(Figure 4C)**. WGA/PI staining of infected barley leaves revealed that DS200-FvRibo1 strain is mostly restricted to the penetration area and rarely reaches the vascular bundles, while DS200 colonized leaf veins and successfully accessed the host vascular bundles at 6 dpi **(Figure 4D)**. In line with this, fungal biomass quantification of the DS200-FvRibo1 and DS200 strains on infected barley leaves at 6 dpi confirmed a significant virulence reduction of the DS200-FvRibo1 compared to DS200 strain **(Figure 4B)**. Together, these findings show that heterologous expression of FvRibo1 attenuated *U. hordei* infection **(Figure 4B-D)**.

**Figure 4.**
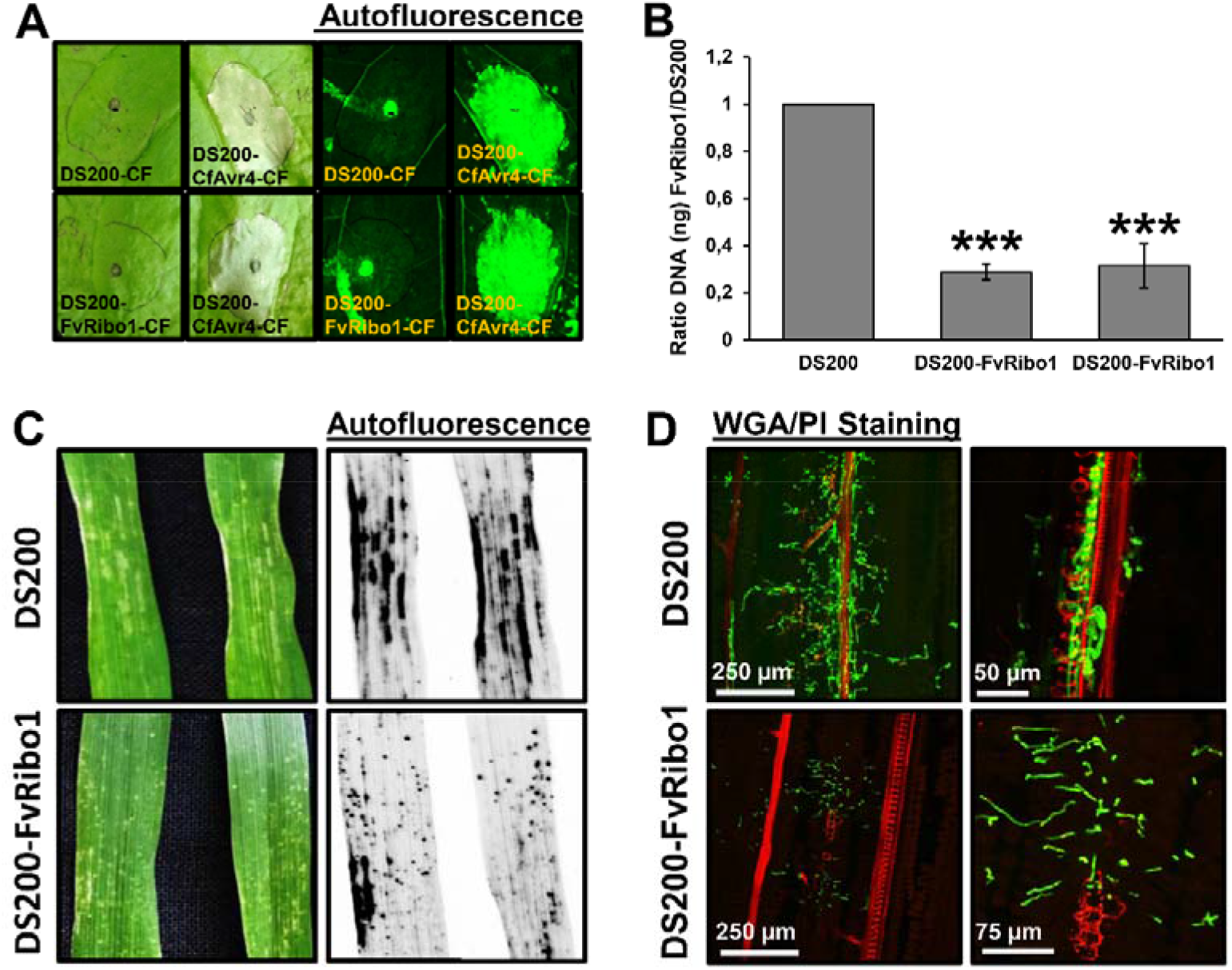
Heterologous expression of fungal effectors in *Ustilago hordei.* **(A) Heterologous expression and secretion of CfAvr4 in** *U. hordei* **DS200 strain** *in vitro*. *U. hordei* strain DS200 expressing *CfAvr4* of *Cladosporium fulvum* with *UHOR_02700* signal peptide and under the control of *pActin* promoter (for constitutive expression), and FvRibo1 of *Fusarium verticillioides* with *UHOR_02700* signal peptide and under the control of *pUHOR_02700* promoter (for expression *in planta* only) were grown in YEPS_light_ liquid medium till OD:1.0. The *U. hordei* cell suspensions were centrifuged and the culture filtrates (CF) of each sample were infiltrated into tobacco leaves expressing Cf4 resistant protein that can recognize CfAvr4 and induce cell death by means of hypersensitive response. The culture filtrates from *U. hordei* DS200 and DS200-FvRibo1 strains were used as negative controls. Pictures were taken at 5 dpi. Autofluorescence of infected leaves was imaged to more easily see sites of cell death by using Gel-Doc (Bio-Rad). **(B) Biomass quantification of DS200-FvRibo1 in barley leaves.** The virulence of the *U. hordei* DS200 and two independent DS200-FvRibo1 strains was assessed by fungal biomass quantification from DNA isolated from infected barley leaves at 6 dpi. The *Ppi1* gene of *U. hordei* was used as a standard for qPCR. The fungal biomass was deduced from a standard curve. A student t-test was performed to determine significant differences, which are indicated as asterisk (***, *P*< 0.001). Error bars represent the standard deviation of three biological repeats. **(C) Heterologous expression and secretion of FvRibo1 in** *U. hordei* **strain DS200** *in planta*. *Ustilago hordei* strain DS200 and DS200 expressing *FvRibo1* (encoding a secreted ribotoxin) of *Fusarium verticillioides* with *UHOR_02700* signal peptide and under the control of the *UHOR_02700* promoter (for only *in planta* expression) were inoculated on susceptible 12-day-old barley seedlings. Macroscopic pictures were taken at 6 dpi. Autofluorescence pictures were taken to see better cell death by using Gel-Doc (Bio-Rad). **(D) WGA-AF488/Propidium iodide staining** was performed to visualize the colonization of DS200-FvRibo1 in barley leaves compared to DS200. While green signal indicates fungal colonization, the red signal represents the plant cell walls.

## Discussion

Plant pathogenic smut fungi are non-obligate biotrophic fungal pathogens, which can infect many economically important crops, such as maize, wheat, barley, oat and sugar cane. Recent comparative genome analysis of five plant pathogenic smut fungi, including *U. hordei, U. maydis*, *Sporisorium reilianum*, *Sporisorium scitamineum* and *Melanopsichum pennsylvanicum*, showed that all of these smut fungi have relatively small genomes (about 20 Mbp) [43]. These genomic features make smut fungi excellent candidates for functional genetic and genomic approaches. The availability of complete genome assemblies, available transcriptomics data and the possibility of performing reverse genetics make the *U. hordei*-barley system a potential model for studying the molecular basis of plant-pathogen interactions. To enable and speed-up functional genetics in the *U. hordei*-barley interaction, we have established several molecular tools, including a solopathogenic *U. hordei* strain, heterologous gene expression and an efficient CRISPR/Cas9 gene editing system.

### Establishment of a solopathogenic strain

The generation of haploid solopathogenic *U. hordei* strains, which express an active bE/bW heterodimer and form infectious filaments without having to fuse with a mating partner, increases the efficiency of genetic transformation by reducing lab workload, since no duplicate mutants in opposite mating partners needed. Quantification of appressoria formation and penetration efficiency of solopathogenic strains showed that there is no significant difference compared to wild-type strain on barley leaves. This result indicates that the solopathogenic strains can be used for functional characterization of virulence factors during barley leaf colonization. Although the barley leaf infection assay did not show any difference in colonization of solopathogenic and wild-type strains, in barley seed infection assays, the solopathogenic strains rarely colonized barley inflorescences and produced teliospores compared to the wild-type strain. This observation indicates that the solopathogenic strain is only weakly pathogenic in systemic colonization of barley and may not produce all virulence factors/effectors needed at the later stages of infection. Accordingly, two generated solopathogenic *U. maydis* strains, SG200 and CL13, also showed attenuated virulence compared to wild-type strains [44, 45]. Moreover, CL13, which is generated by replacement of only compatible *b* loci, shows a more attenuated virulence compared to the SG200 strain, which is generated by replacement of both compatible *a* and *b* loci [44, 45]. In our experiments, the *U. hordei* solopathogenic strains DS199 (with only compatible *b* alleles) and DS200 (with both compatible *a* and *b* alleles) showed similar rates of virulence during barley penetration. This suggests that the presence of a compatible *a* loci is more important for the formation of infectious filaments in *U. maydis*, which has a tetrapolar mating system, than in *U. hordei*, which has a bipolar mating system [19].

Recently, Schuster *et al*., (2016) established a CRISPR/Cas9 gene editing system for *U. maydis* with ~70% efficiency [46]. By using a similar approach, we achieved a very efficient CRISPR/Cas9-HF based system in *U. hordei* with ~83% gene editing efficiency in progeny. CRISPR/Cas9 gene editing of the *U. hordei Fly1* gene, a fungalysin metalloprotease involved in fungal cell separation [41], resulted in an impaired cell separation phenotype in DS200, indicating the functional conservation of this protein among smut fungi. Thus, establishment of both solopathogenic strains and a CRISPR/Cas9 gene editing system allow fast and efficient reverse genetic approaches in *U. hordei*.

### Ultrastructural analysis of the *Ustilago hordei*-barley biotrophic interphase

During barley leaf colonization, *U. hordei* enters in the host cell without breaching the host plasma membrane. Thus, the *U. hordei*-barley interaction is mediated through the biotrophic interphase, which comprises the fungal cell wall (FCW), electron-dense and electron-translucent extracellular matrixes (edECM and etECM) and the plant cell wall (PCW) (in some part of colonized tissues). Presence of vesicles in the fungal cytoplasm close to the hyphal tip and in the surrounding plant cytoplasm as well as of plant multivesicular bodies close to fungal penetration sites indicates that the biotrophic interphase is very active site. Some vesicles appeared to be in the process of either fusing with or pinching off the plant cell membrane. The edECM of unknown composition surrounds the FCW and in some regions, its outer surface shows irregular patterns with small protrusions. At contact sites with the PCW and at cell-to-cell penetration sites, *U. hordei* accumulates a thicker edECM and it seems that the material causing the electron-density of the ECM diffuses into the adjacent PCW. This observation indicates that the interphase between the plant cell membrane and the edECM is quite active. The plant apoplastic space contains a wide range of defense components, such as glycoside hydrolyses, proteases, peroxidases, antimicrobial proteins and secondary metabolites, which collectively contribute to plant immunity [47]. *U. hordei* hyphae may secrete the edECM to prevent access of these plant-derived defense components to the fungal cell. Since the thickness of edECM gets higher with contact to the host PCW, one can also hypothesize that it is required for anchoring to the PCW to increase cell-to-cell penetration efficiency. Similar edECM were also observed for other smut fungi, such as the maize smut fugus *U. maydis* and *Ustacystis waldsteiniae* (on *Waldsteinia geoides* host), rust as well as powdery mildew (*Hyaloperonospora parasitica*) during host colonization [48–51]. Although the content of edECM is unknown, immunogold gold labeling with (1-3)-β-glucan specific antibodies revealed presence of callose in the electron-translucent extracellular matrix (etECM). In incompatible host-pathogen interactions, callose deposition at the site of infection is a hallmark of plant immune response; however, in compatible interactions, successful pathogens (like *U. hordei*) can suppress the host defense response, including callose deposition [52]. Therefore, the presence of callose in the etECM indicates that *U. hordei* may have an ability to use callose at the biotrophic interphase as an additional carbon source. Up-regulation of several 1,3 beta-glucanase encoding genes in *U. hordei* during host colonization might be involved in this process [53].

To reach the host vascular bundles, *U. hordei* grows/moves mostly intracellularly from cell-to-cell. Detailed observation revealed that at the site of cell-to-cell penetration, the *U. hordei* hypha developed a swollen structure, which resembled an appressorium. A similar phenotype of swollen hyphal tips was also observed in *U. maydis* during cell-to-cell penetration in maize [54]. Formation of this structure may increase the penetration efficiency of smut fungi from cell-to-cell movements. In Ökmen *et al*., (2018), we have reported that *U. hordei* intracellular hyphae develop lobed haustoria-like structures during barley colonization [20]. One infected host cell could have more than one haustorium. Haustorial structures of *U. hordei* were distinguished from the normal hyphae by their bigger size and inter-connected lobular shapes. While both rust and powdery mildew haustoria are formed from extracellular fungal hyphae [55, 56], *U. hordei* haustoria structures originate from intracellular hyphae. In addition, a structure comparable to the neckband of rust haustoria that separates the haustorial matrix from the apoplast was not seen in our analysis [57]. Detailed transmission electron micrographs of *U. hordei* haustoria also showed that these structures contain large vacuoles with a fine-granular content and intraluminal vesicles.

### Heterologous expression of fungal effectors in *Ustilago hordei*

Both smuts and rusts are plant pathogenic fungi belonging to the division Basidiomycotina [58]. In addition to their phylogenetic relationships, smuts and rusts (obligate biotroph) have similar biotrophic life styles, in which they require an intimate association with their hosts to acquire nutrients and complete their pathogenic lifecycles. Apart from their phylogenetic relationships and similar lifestyles, *U. hordei*, *Bgh* and *Pgt* also form comparable intracellular haustorial structures, secrete edECM, and infect the same host plant species. Due to host-specific adaptations during co-evolution, entirely different pathogens that share the same host can independently develop different types of effectors that interact with the same target. Accordingly, Avr2 of *C. fulvum* (fungus) [59–61], EPIC1 and EPIC2B of *Phytophthora infestans* (oomycete) [62], Gr-VAP1 of *Globodera rostochiensis* (nematode) [63] and Cip1 from *Pseudomonas syringae* (bacterium) [64] can interact and inhibit tomato cysteine protease, Rcr3. Therefore, the *U. hordei*-barley pathosystem allows to perform reverse genetics for *Bgh* and *Pgt* effectors in the same host system and subsequently to identify their virulence functions.

Successful heterologous expression and secretion of the GUS-mCherry recombinant protein in *U. hordei* during barley colonization demonstrates that the designed concept is feasible. Some of the GUS-mCherry was also cleaved in apoplastic space of barley leaf, indicating C-terminal processing of this protein. For further proof of concept that *U. hordei* can express and secrete functional effectors from different fungi, a very robust Avr4/Cf4 pair was used to induce HR-mediated cell death. To this end, *Avr4* avirulence gene from *C. fulvum* was expressed in DS200 *in vitro*. Induction of Cf4-mediated HR only in the presence of culture filtrate of DS200-Avr4 strain indicates that DS200-Avr4 expresses and secretes functional Avr4 from a biotrophic *C. fulvum* that can be recognized by Cf4 resistance protein. In a similar way, *in planta* expression and secretion of a plant cytotoxic *Ribo1* (FvRibo1) from *F. verticillioides* in DS200 was confirmed by using only *in planta* expressed *U. hordei* promoter. Heterologous expression of the FvRibo1 in DS200 negatively affects the colonization of this biotrophic smut fungus in barley. The WGA-AF488/PI staining and biomass quantification assays display that DS200-FvRibo1 hardly moves and colonizes vascular bundles and showed significantly reduced fungal biomass compared to DS200 strain. Moreover, macroscopic and microscopic observations of focal necrotic spots on barley leaves indicate that FvRibo1 is also cytotoxic to barley cells. Heterologous expression of exogenous effector genes from different plant pathogenic fungi showed that the solopathogenic *U. hordei* DS200 strain can be used for functional characterization of effectors from biotrophic phytopathogens as well.

Biotrophic infection of barley leaves, formation of intracellular hyphae and haustorial structures as well as the molecular tools presented in this study make the *U. hordei* pathosystem as a useful platform for the functional analysis of effector proteins from biotrophic fungi.

## Supporting information

Supplementary Figures with legends

## Acknowledgements

We thank Ila Rouhara for technical assistance for TEM. We acknowledge Regine Kahmann and the Max-Planck-Institute for Terrestrial Microbiology, Marburg, Germany, for providing generous support and access to infrastructure. This research is supported by the European Research Council under the European Union’s Horizon 2020 research and innovation program (consolidator grant conVIRgens, ID 771035).

## Author contributions

BÖ and GD designed the research; BÖ, UN and DS performed experimental work, BÖ analyzed the data. GB provided materials and advice on the experimental design, BÖ wrote the paper with input from GD, UN and GB.

## Conflicts of Interest

The authors have no conflicts of interest

## Supplementary Material

**Figure S1** *Ustilago hordei-***barley infection assay.** Dehulled and surface sterilized barley Golden Promise seeds were inoculated with the *U. hordei* wild-type and DS200 strains. Approximately 3-4 months after inoculation, disease symptoms were observed at barley heading.

**Figure S2 (A-D) Cell-to-cell penetration of** *Ustilago hordei*. *U. hordei* primarily grows intracellularly at 8 dpi in barley leaves. The fungal hyphae grow through the plant cells and frequently branch. The *U. hordei* hyphal cell wall was surrounded by an extracellular matrix of electron-dense (edECM) (between 50-500 nm thickness) of unknown composition. The extracellular matrix gets thicker where the hypha was in contact with the plant cell wall (white arrowheads in **A**). edECM: electron-dense extracellular matrix; LB: Lipid bodies; PCW: Plant Cell Wall.

**Figure S3 (A-C) Ultrastructural features of the** *Ustilago hordei* **during barley colonization.** The *U. hordei* electron-dense extracellular matrix (edECM) gets thicker where the hypha was in contact with the plant cell wall (PCW) and has a highly irregular surface structure **(C)**. **(D-F)** *U. hordei* hyphae contain free ribosomes, strands of endoplasmic reticulum (ER), mitochondria (m), nuclei (n), as well as lipid bodies (LB), vesicles (Ve) and vacuoles (Va). Transmission electron micrographs also showed the presence of closely paired nuclei, tightly associated with mitochondria **(F)**. Vesicles (Ve) with cores of different electron densities and multivesicular bodies (MVB) were detected in hyphal tips and in the plant cytoplasm close to fungal penetration sites, respectively **(B, C, F)**.

**Table S1.** Plasmids and primers used in this study.

## Notes

### Competing Interest Statement

The authors have declared no competing interest.

