## Supplementary Figures with legends for "The *Ustilago hordei*-barley interaction is a versatile system to characterize fungal effectors"

### Supplementary Material

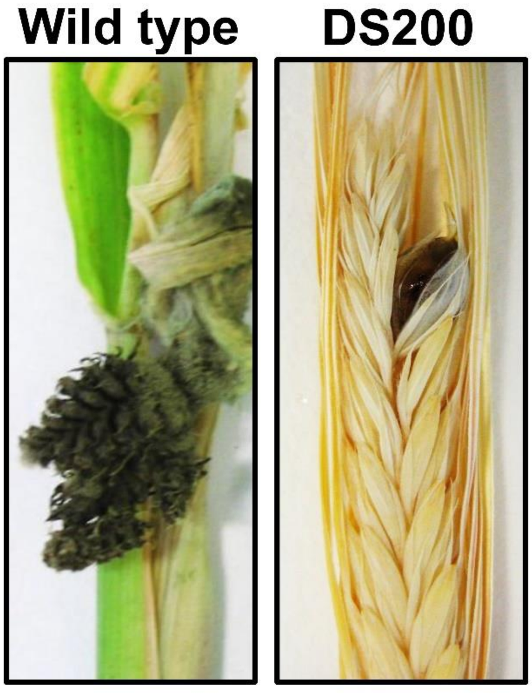


**Figure S1 *Ustilago hordei*-barley infection assay.** Dehulled and surface sterilized barley Golden Promise seeds were inoculated with the *U. hordei* wild-type and DS200 strains. Approximately 3-4 months after inoculation, disease symptoms were observed at barley heading.

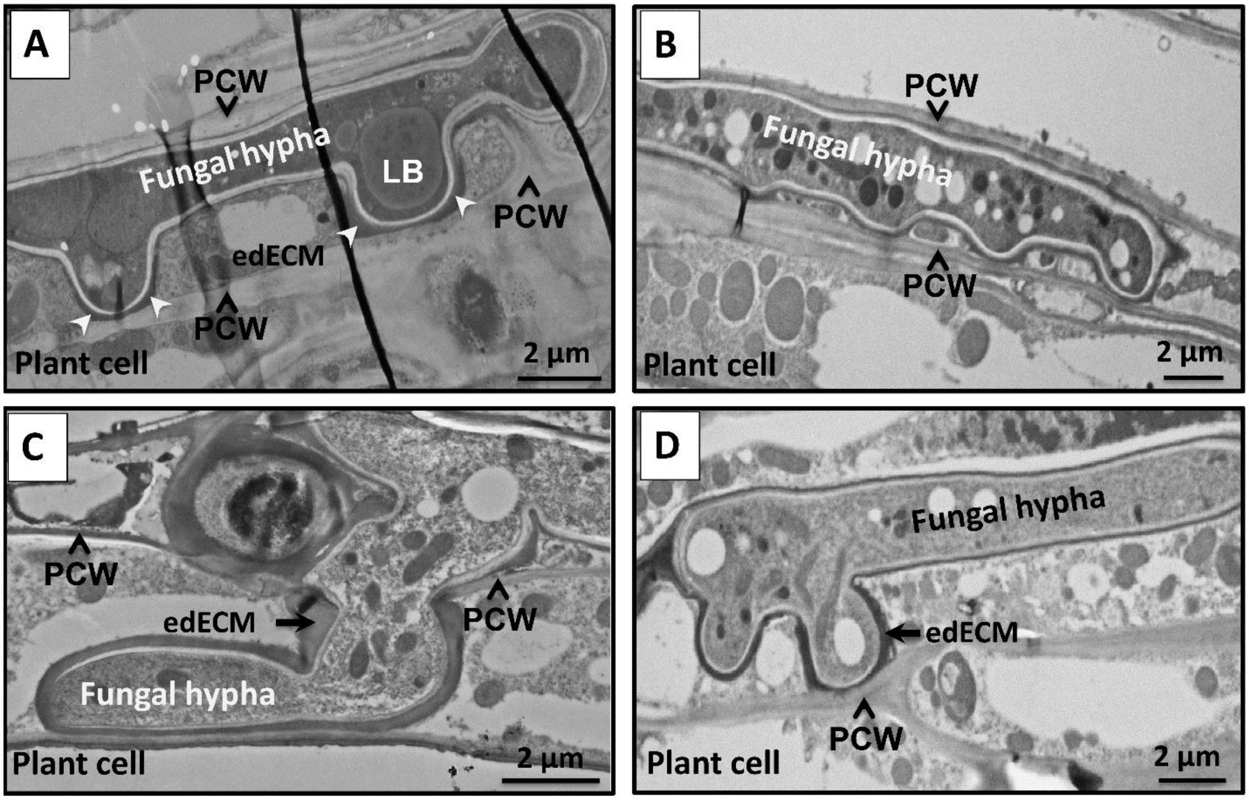


**Figure S2 (A-D) Cell-to-cell penetration of *Ustilago hordei*.** *U. hordei* primarily grows intracellularly at 8 dpi in barley leaves. The fungal hyphae grow through the plant cells and frequently branch. The *U. hordei* hyphal cell wall was surrounded by an extracellular matrix of electron-dense (edECM) (between 50-500 nm thickness) of unknown composition. The extracellular matrix gets thicker where the hypha was in contact with the plant cell wall (white arrowheads in **A**). edECM: electron-dense extracellular matrix; LB: Lipid bodies; PCW: Plant Cell Wall.

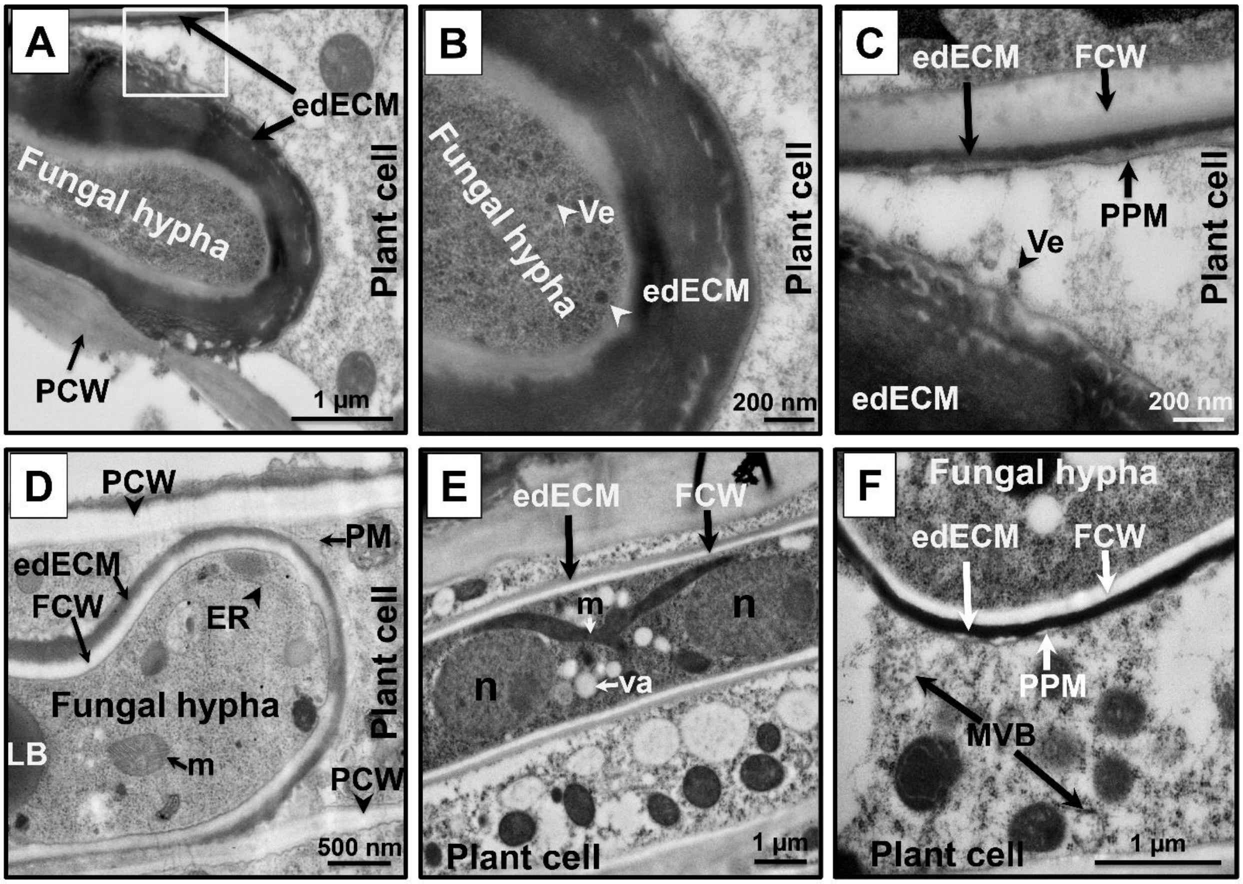


**Figure S3 (A-C) Ultrastructural features of the *Ustilago hordei* during barley colonization.** The *U. hordei* electron-dense extracellular matrix (edECM) gets thicker where the hypha was in contact with the plant cell wall (PCW) and has a highly irregular surface structure **(C)**. **(D-F)** *U. hordei* hyphae contain free ribosomes, strands of endoplasmic reticulum (ER), mitochondria (m), nuclei (n), as well as lipid bodies (LB), vesicles (Ve) and vacuoles (Va). Transmission electron micrographs also showed the presence of closely paired nuclei, tightly associated with mitochondria **(F)**. Vesicles (Ve) with cores of different electron densities and multivesicular bodies (MVB) were detected in hyphal tips and in the plant cytoplasm close to fungal penetration sites, respectively **(B, C, F)**.

**Table S1.** Plasmids and primers used in this study.
